# Lost in Translation: population genomics and long-read sequencing reveals relaxation of concerted evolution of the ribosomal DNA cistron

**DOI:** 10.1101/811216

**Authors:** Keaton Tremble, Laura M. Suz, Bryn T.M. Dentinger

## Abstract

Concerted evolution of the ribosomal DNA array has been studied in numerous eukaryotic taxa, yet is still poorly understood. rDNA genes are repeated dozens to hundreds of times in the eukaryotic genome [1] and it is believed that these arrays are homogenized through concerted evolution [2, 3] preventing the accumulation of intragenomic, and intraspecific, variation. However, numerous studies have reported rampant intragenomic and intraspecific variation in the rDNA array [4–10], contradicting our current understanding of concerted evolution. The internal transcribed spacers (ITS) of the rDNA cistron are the most commonly used DNA barcoding region in *Fungi* [11], and rely on concerted evolution to homogenize the rDNA array leading to a “barcode gap” [12]. Here we show that in *Boletus edulis* Bull., ITS intragenomic variation persists at low allele frequencies throughout the rDNA array, this variation does not correlate with genomic relatedness between populations, and rDNA genes may not evolve in a strictly concerted fashion despite the presence of unequal recombination and gene conversion. Under normal assumptions, heterozygous positions found in ITS sequences represent hybridization between populations, yet through allelic mapping of the rDNA array we found numerous heterozygous alleles to be stochastically introgressed throughout, presenting a dishonest signal of gene flow. Moreover, despite the signal of gene flow in ITS, our organisms were highly inbred, indicating a disconnect between true gene flow and barcoding signals. In addition, we show that while the mechanisms of concerted evolution are ongoing in pseudo-heterozygous individuals, they are not fully homogenizing the ITS array. Concerted evolution of the rDNA array may insufficiently homogenize the ITS gene, allowing for misleading signals of gene flow to persist, vastly complicating the use of the ITS locus for DNA barcoding in *Fungi*.

## Introduction

Concerted evolution, the homogenization of high copy-number repetitive genes, has been widely studied across many eukaryotic taxa and its effects on ribosomal DNA arrays (rDNA) have been extensively observed (Fig: 1)[1]. However, the mechanisms underlying concerted evolution are still poorly understood. rDNA genes are repeated dozens to hundreds of times in the eukaryotic genome and are organized in one to several large tandem arrays [1]. Despite this high copy number that could lead to the formation of divergent paralogs through random mutation, it is believed that these arrays are homogenized through gene conversion and unequal recombination [2, 3] preventing the accumulation of intragenomic, and subsequently, intraspecific variation (Fig. 1). Unequal recombination (UR), a form of homologous recombination among paralogs in the rDNA array, is the most empirically supported mechanism of concerted evolution [4, 5, 13]. However, previous work has only indirectly observed the expected results of UR, and direct support has never been provided. Moreover, gene conversion (GC) in tandem arrays has little empirical support outside of protein-coding gene families [14]. Although intragenomic variation should not persist due to concerted evolution, numerous studies have reported the presence of problematic intraspecific variation in the non-coding regions, particularly the internal transcribed spacers (ITS), of the rDNA array [4–10]. If this intraspecific variation is due to relaxed concerted evolution, our assumptions about the origin of this variation that underpin its application for species delineation need to be reconsidered.

**Figure 1:**
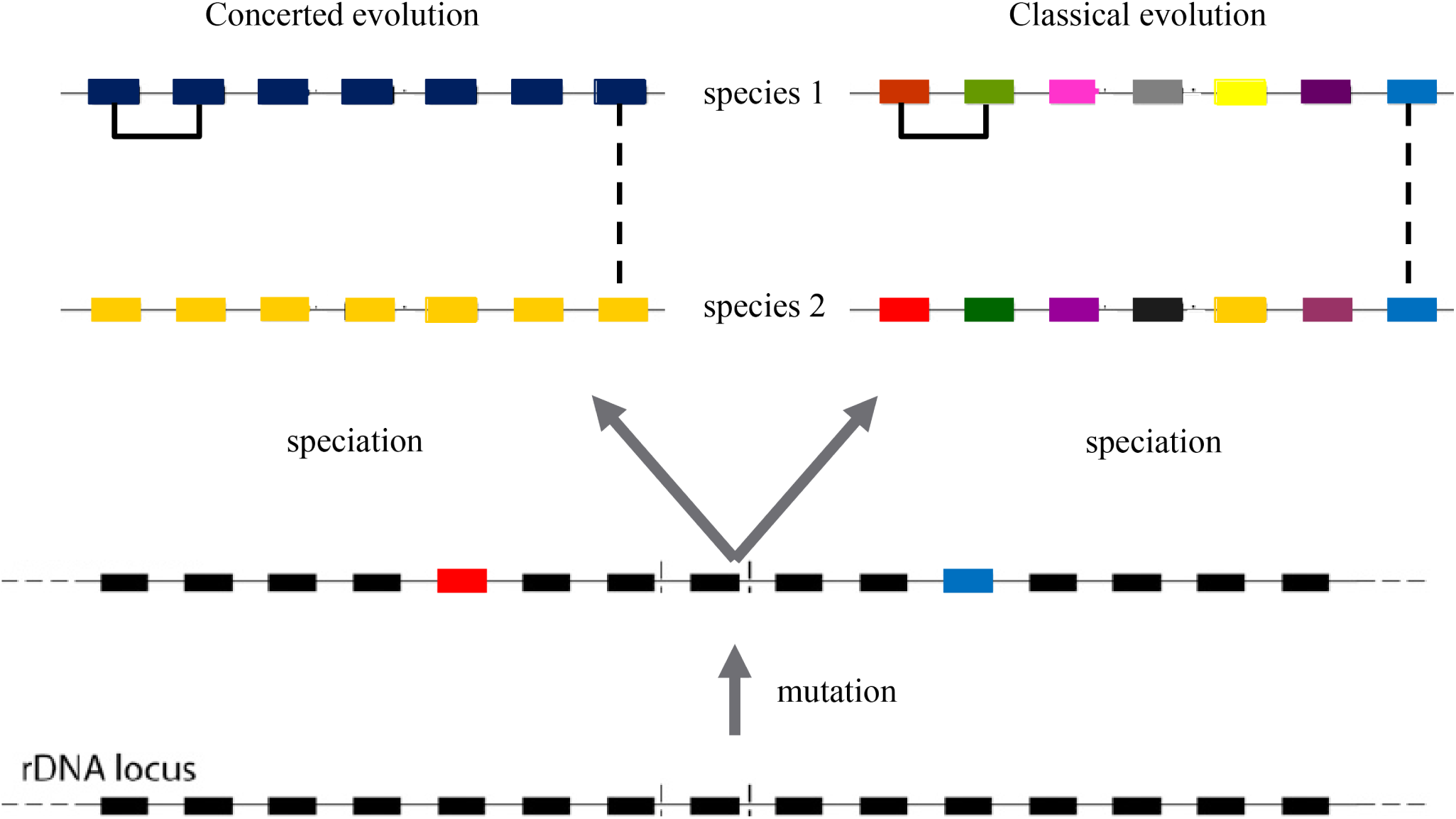
Adapted from Ganley and Kobayashi 2007. The rDNA array has been proposed to evolve in a strictly concerted fashion, preventing accumulation of intraspecific variation; where paralogs (solid black lines) have more similarity than orthologs (dashed lines). This is in contrast to gene evolution via classical evolution; where orthologs share more similarity than paralogs.

The internal transcribed spacers (ITS) of the rDNA cistron, which are transcribed but not translated, are the most widely used DNA barcoding region in *Fungi* [11] with far-reaching impacts across many scientific disciplines. For example, according to a Web of Science search on 12 September 2019, in the last five years, there were 12,822 journal articles published with the words “fungi” or “fungal” or “fungus” and “internal transcribed spacer” in the title. A little more than half (55%) of all newly identified fungal species from 2011-2016 were deposited with ITS sequences [15]. More-over, the ITS region is the predominant tool for linking new fungal names to type specimens [16]. The power of ITS barcoding lies in the rapid accumulation of polymorphisms between species, with limited intraspecific variation, leading to a “barcode gap” [12]. Concerted evolution is thought to be the primary mechanism preventing accumulation of intraspecific variation in ITS, yet a “relaxation” of this homogenizing force has been found in numerous taxa, yielding significant ITS variation [5, 10]. This variation can be the manifestation of reproductive isolation when corroborated by independently inherited loci. However, a growing body of research has documented ITS variation found within even a single individual. For example, ITS-PCR cloning from multiple *Amanita* cf. *lavendula* specimens revealed a diversity of sequences with an average sequence similarity of 96.93% [10]. More examples of significant intragenomic and intraspecific variation have been found across the *Fungi*, but whether this variation persists within or between cellular genomes is unknown [5, 8, 17].

Due to the large number of nearly identical tandem rDNA repeats that prohibits accurate recon-struction from short-read DNA sequence data, little is known about the structure of this documented ITS variation within the collective genomic rDNA array. Yet, the location of ITS variation within the genome can have important biological implications. For example, a heterozygous position in ITS could be interpreted as hybridization and ongoing gene flow between populations, or incomplete lineage sorting from recent reproductive isolation. However, these interpretations may be incorrect if the heterozygous alleles are introgressed within tandem arrays rather than between parental chromosomes, indicating instead mechanisms independent of recent gene flow and recombination. PCR cloning followed by Sanger sequencing allows identification of variation within the collective genome of an individual, but provides no information concerning the location of any one sequence in relation to other alleles. In addition, Sanger sequencing produces a consensus sequence that captures only the information in highest frequency along the sequence [6]. High-throughput sequencing technologies, such as Illumina’s sequencing-by-synthesis, offers greater resolution due to sequencing of individual DNA fragments, but approaches to identify intragenomic variation from these data have also been problematic due to the need to reconstruct repeated elements from overlapping short sequences that do not span the full length of the repeat. Due to the highly repetitive nature of rDNA genes, traditional bioinformatic approaches that attempt to assemble contiguous sequences from overlapping short reads collapse the arrays into a single contiguous sequence and only retain polymorphic information if it exists in Hardy-Weinberg equilibrium. Moreover, amplicon-based approaches such as those that dominate metagenomics profiling of microbiome and environmental samples, will similarly capture only the polymorphisms at high frequency, potentially underrepresenting variation that could lead to spurious taxonomic assignment. While high-throughput genomic sequencing continues to revolutionize the field of phylogenetics, current bioinformatic tools are inadequate for accurately capturing the full variation in the primary barcoding region using in *Fungi*.

Large, widespread species are ideal targets for investigating patterns of intragenomic ITS variation because they are more likely to harbor polymorphisms than smaller populations with limited ranges and present a challenge to taxonomic delineations using DNA barcodes. The core porcini mushroom species *Boletus edulis* Bull. is a globally distributed and economically important wild, edible ectomycorrhizal mutualist in which we have observed from a dataset of >200 samples rampant intraspecific ITS variation, including highly structured populations and extensive heterozygosity [18, 19]. Sequence similarity in the dataset ranged from 0.812 to 1.0. Importantly, the independently inherited translation-elongation factor 1-alpha (EF1-alpha) single copy gene sequences from *B. edulis* do not corroborate this intraspecific diversity found in the ITS (Fig. 2), indicating this variation cannot be strictly interpreted as indicating gene flow within and between populations. Therefore, this ITS diversity may present significant challenges for taxonomic delineations using traditional DNA barcoding methods. For example, the current standard 1.5% sequence similarity species cutoff [20], appears to be thoroughly inappropriate when sequences within a single individual may differ by approx. 10%. While some of the taxonomic shortcomings of ITS have been previously highlighted [15], the mechanistic cause behind the persistence of intraspecific variation is entirely unknown and has simply been thought to be a relaxation of concerted evolution [1]. To preserve the continued use of the ITS region as a taxonomic barcode it is imperative that we understand the processes behind the creation of intragenomic variation within the context of concerted evolution, and how these variants give rise to the patterns observed at the population and species levels of organization.

**Figure 2:**
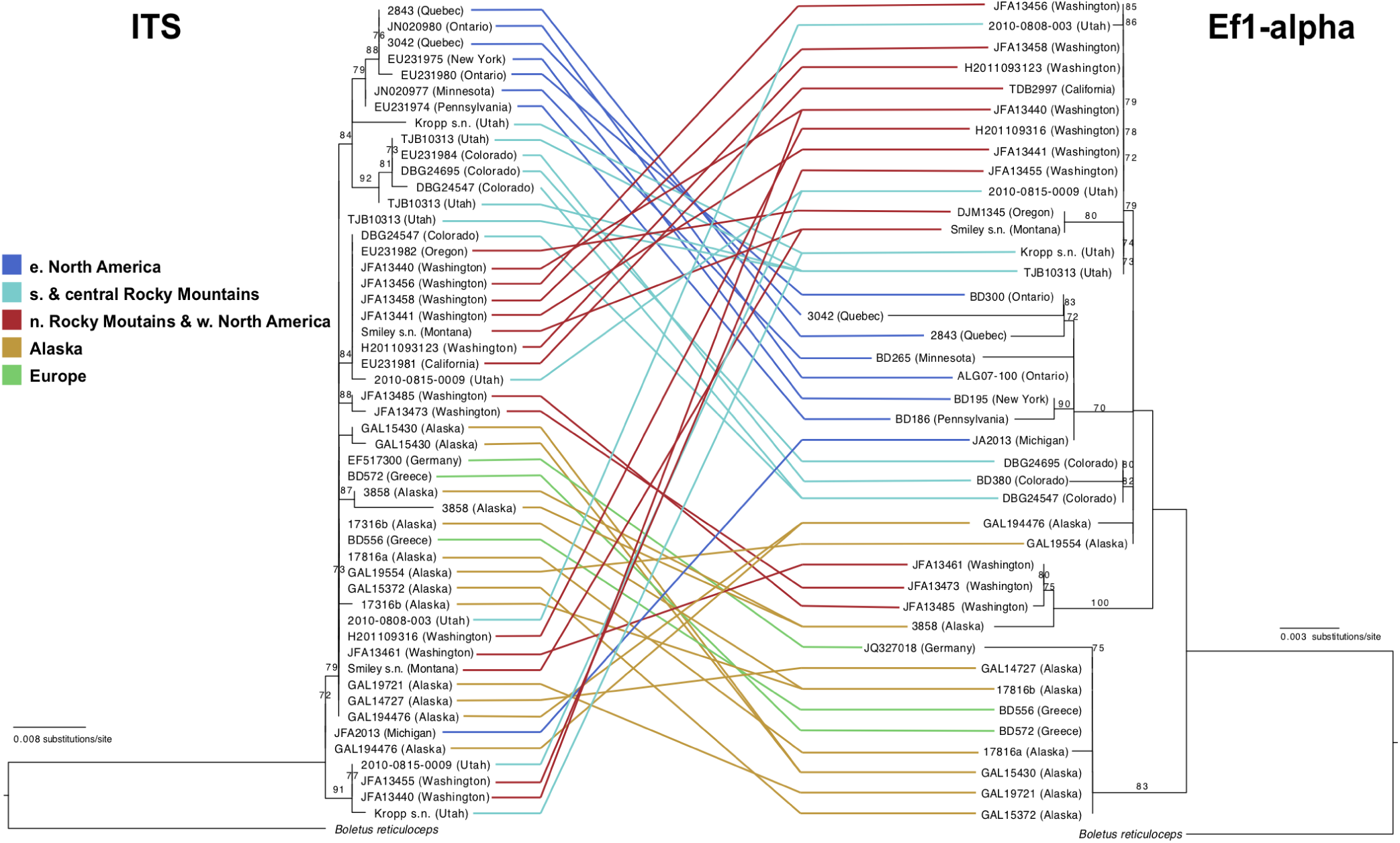
Comparison of *B. edulis* ITS (left) and EF1-alpha (right) maximum-likelihood phylogenies. The ITS dataset consists of phased polymorphisms observable in Sanger sequencing trace files and confirmed by PCR-cloning. Sequences were aligned with the L-INS-i algorithm in MAFFT. ModelFinder was used in IQ-TREE to find the best partitioning scheme (ITS partitions=ITS1,5.8S,ITS2; EF1-alpha partitions=exonic 1^*st*^,2^*nd*^,3^*rd*^ codon positions and introns) and model (option MF+MERGE), allowing each partition to have its own evolutionary rate (option-spp). Branch support was assessed using 1000 ultrafast bootstraps (option -bb) and resampling partitions and then sites within resampled partitions (option -bpsec GENESITE). From this comparison we highlight the discrepancy between relatedness in ITS sequences and other barcoding loci.

To identify ITS variation in *B. edulis*, and understand how variation persists despite concerted evolution, we developed new methods to manually reconstruct ITS consensus sequences from 29 high- and low-coverage whole genome sequences. We then compared these genomic ITS sequences to sequences produced through Sanger sequencing to assess the impact of using different sequencing methods to highlight ITS variation and the consequences for phylogenetic-based species recognition. Finally, to map the allelic structure of the ITS array for the first time in any organism, we used Oxford Nanopore Technology’s nanopore sequencing (ONT) to generate continuous long reads (>10kb) that spanned multiple rDNA cistrons.

## 1 Materials and Methods

### 1.1 High-Throughput Sequencing

Twenty-seven of the 29 samples were obtained from dried specimens (Table S1) or from samples stored in ethanol provided by Dr. Amos of the University of Cambridge. Genomic DNA from dried specimens collected in England (all samples besides BD747 and MO278521) was extracted following a CTAB protocol [21]. DNA from specimens stored in alcohol and previously dried for an estended period of time, was carried out using a QIAGEN DNeasy Plant mini kit. Libraries from specimens were prepared using a TruSeq Nano DNA LT (Illumina Inc.) sample kit with a final insert size of 550 bp. The ten indexed libraries were normalized and pooled based on an approximately 30-fold depth per sample. Paired-end sequencing (2 × 300 bp) was performed in an Illumina MiSeq sequencer in the Jodrell Laboratory at the Royal Botanic Gardens, Kew. In addition, two new samples, assumed to be heterozygotes, BD747 from Utah and MO278512 from Connecticut, were extracted and sequenced separately. Total DNA was extracted with the Zymo PowerSoil DNA extraction kit according to the manufacturer’s protocol and sequenced on 2 × 150 PE Illumina HiSeq2500 by RAPiD Genomics (Gainesville, FL). In addition, a high molecular weight DNA library was prepared for BD747 using a SDS-based lysis buffer and phenol:chloroform extraction and the one-pot ligation protocol [22]. This library was sequenced using one R9.5 flow cell on an Oxford Nanopore Technology MinION.

### 1.2 Sanger Sequencing

DNA for amplification was extracted in KCl-tris-HCl at 95 degrees C for 10 minutes, and then stored with an equal volume of a 3% BSA solution. The ITS region was amplified using the Agaricomycete primers ITS8F and ITS6R and the corresponding PCR protocol outlined in [21]. Amplification success was verified by gel-electrophoresis and Sanger sequencing was performed at the DNA Sequencing Core Facility, University of Utah.

### 1.3 ITS contig construction and heterozygote identification

After adapter removal with with fastp (v0.20.0) [23] ITS sequences were reconstructed from 28 *B. edulis* genomes in two ways 7: 1) using standard bioinformatic methods and 2) using a novel raw-read reconstruction pipeline. Using method 1, trimmed genomes were aligned to a reference genome assembly, produced from BD747, with Bowtie2 (v2.3.5.1)[24]. Aligned reads were assembled with Spades (v3.11.1) [25]. In addition, a hybrid assembly of BD747 from Illumina and Nanopore reads was produced with MaSuRCA [26]. ITS sequences were recovered from assemblies using BLASTn with an ITS reference from BD747 used as a query. Through method 2, raw Illumina sequences from each genomes were aligned directly to the B. edulis ITS reference with Bowtie2, assembled with Mafft (V7.0) [27] using the FFT-NS-2 algorithm, and a consensus sequence was constructed using a conservative 15% frequency cutoff for inclusion in the consensus. Heterozygous alleles were found to exist at lower frequencies, however, lower cutoffs produced many false-positives at low coverage sites and were prohibitively time consuming to correct. To calculate heterozygous allele frequencies at each position, a search through the raw read files, including four nucleotides on either side of the polymorphism, were recorded along with the number of raw reads matching either allele. After construction of consensus sequences, 28 contiguous ITS sequences produced from Illumina reads were aligned with the Mafft L-NS-i algorithm, and any polymorphism in the alignment between the sequences was checked for validity and heterozygosity from the raw reads. The absolute cutoff for determining “true” reads (i.e. heterozygous alleles present in greater numbers than would be expected from error) was 0.5% of total read count at the position and a minimum of 5 corroborating reads. We consider this to be a conservative cutoff because it is twice as stringent as any reported error for Illumina single-nucleotide substitutions [28]. All statistics were performed in Rstudio Version 1.2.1335

### 1.4 Genomic derived ITS and Sanger sequenced ITS comparison

Five genomic and Sanger derived ITS sequences, from BD747, BD572, 20100815009, MO278512, and DBG24695, were aligned and all polymorphic positions were assessed. Specifically, any polymorphic region found in the genomic ITS sequences was checked for presence or absence in the corresponding Sanger sequence.

### 1.5 Identification of allelic structure of rDNA array

The primary constraint with any analysis of the rDNA tandem array is the innate highly repetitive structure. This prevents the localization of any read to a single ITS region or even array. Because reads cannot align separately to either array, heterozygous alleles may be present uniformly within one array as would be the case in a F_1_ hybrid, or introgressed into both arrays via some mechanism of concerted evolution. Without further information both cases may be equally likely when analyzing genomic ITS sequences. To overcome this repeat-rich region we sequenced one specimen (BD747) with the MinION long read sequencer from Oxford Nanopore Technology. The raw MinION reads were then aligned to our *B. edulis* reference, and previously identified heterozygous positions were searched for among the raw reads. Due to the reportedly high error rate associated with Nanopore Sequencing, no MinION reads were used to identify novel alleles, only to confirm positioning along the ITS array. We performed three separate allele search regimes to characterize allelic structure outlined below (Fig. 4C).

**Figure 3:**
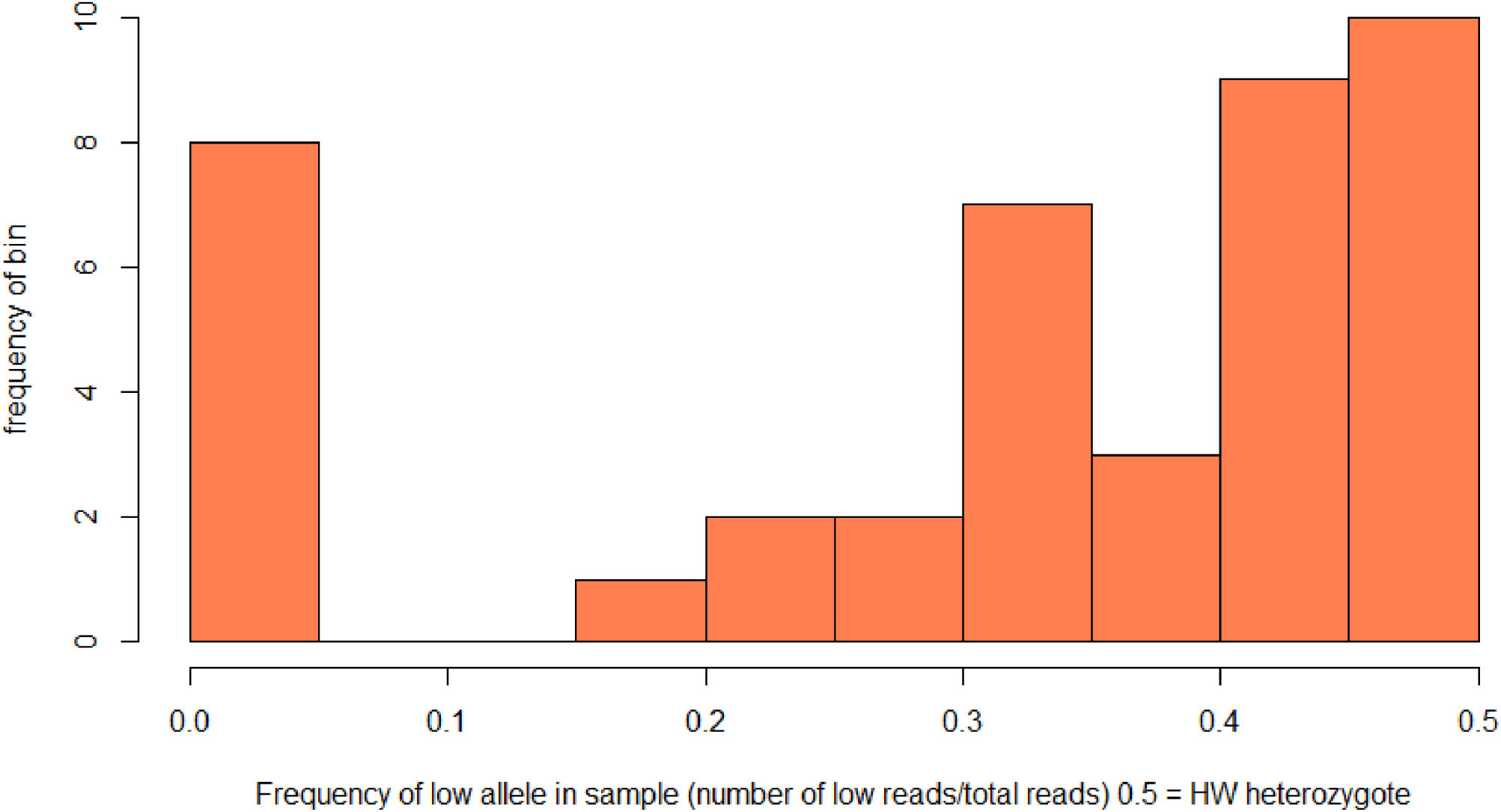
Histogram of ITS allele frequencies at heterozygous positions produced from raw Illumina read alignments. A value of 0.5 on the horizontal axis represents a 50/50 hybrid where the number of reads matching each allele were approximately equal.

**Figure 4:**
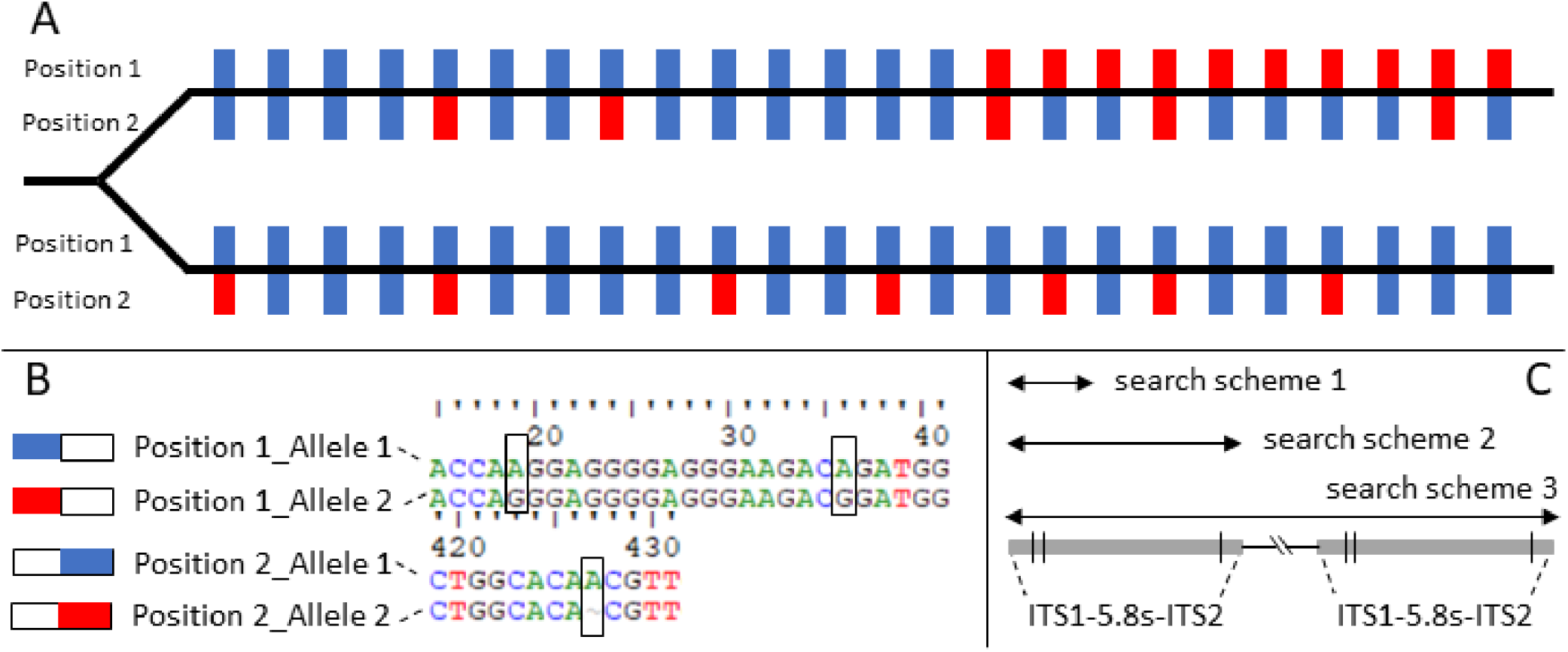
A) Approximated allelic structure of the ITS tandem array in BD747 revealed through ONT long read sequencing. Blue color indicates that the ITS region possess the majority allele at that heterozygous position while red indicates the minority allele. ONT reads spanning the entire ITS were used to link the first heterozygous position with the second, and ultra-long reads spanning two ITS regions were used to characterize allelic structure. If alleles were found in two consecutive reads, they were assumed to be located within the same array. B) Allele 1 and 2 at position 1 had no introgressed long or short reads, indicating that they persist as separate blocks. C) Raw read search schemes used to identify ITS haplotype diversity and array allelic structure. Black lines in the ITS regions represent the position of the three heterozygous alleles in part B. Search 1 identified short reads that covered the first two alleles at position 1 (approx. 50bp window); search 2 identified MinION reads that covered position 1 and 2 in the same ITS gene (approx 680bp window); search 3 identified MinIOn reads that covered position 1 and 2 for two consecutive ITS genes (approx 10kb window). In each search scheme, the number of reads presenting each combination of alleles was counted to estimate haplotype abundance and approximate location found in A.

**Search 1** We searched for raw Illumina reads that covered the first two A/G heterozygous positions of BD747 (approx 50bp window)(Table S2). Both positions posses a majority allele (position 1: A—97.2% of reads, position 2: A—78.2% of reads) and a minority allele (position 1: G—2.8% of reads, position 2: G—21.8% of reads). Searches were performed to identify reads that contained alleles in all combinations (i.e. position 1: A – position 2: A, position 1: A – position 2: G).

**Search 2** Using raw BD747 MinION long-reads, we searched for reads that spanned the entire ITS region. The length of the ITS region necessitates the use of long read sequencing. From these reads, we identified reads that covered the first heterozygous position (A/G) and the third heterozygous position 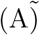 (approx 680bp window)(table S2) in all allele combinations.

**Search 3** To identify the positional relationships of any two ITS regions, ultra-long (>10kb) MinION reads that spanned multiple rDNA cistrons were used. From these ultra-long reads, we searched for the majority and minority alleles of heterozygous positions 1, 2, and 3, and identified the number of reads that presented each combination of haplotypes. For example, we searched for reads that contained A – A in the first cistron and G – G in the second. If two haplotypes, each representing separate ITS genes, were found within the same ultra-long read, this indicates that they are located within the same rDNA array.

### 1.6 ITS Dosage Calculations

Unequal recombination, during the process of concerted evolution, has been show to shift rDNA copy numbers [5]. To approximate rDNA copy number and potentially identify the occurrence of unequal recombination in our specimens, we estimated rDNA dosage according to [29]. While not a precise estimate of rDNA copy number, dosage calculation may allow us to highlight the magnitude of unequal recombination and copy number diversity in our specimens.

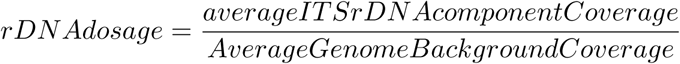

After alignment to BD747 with Bowtie2, Samtools (v1.4)[30] depth was used to calculate average sequence depth at each position for both the ITS region and the whole genome. To determine if the variance in dosage was a byproduct of sequencing depth where high coverage sequencing runs produced disproportionately high ITS coverage, we ran a linear regression model and found no significant relationship (p = 0.84) between the variables. In addition, the program BUSCO (V3) with the Basidiomycota training set was used to characterize genomic sequence quality [31]. As a secondary measure of sequencing success, the number of missing BUSCO elements, was used to asses rDNA dosage and no significant relationship was found.

### 1.7 Whole genome variant calling, inbreeding estimation, and SNP haplotyping

Variants were called with GatK version (4.2.1.0) [32] using BD747 as a reference, according to their prescribed best practices with several notable exceptions: 1) to recalibrate raw base scores, three rounds of SNP calling were conducted and hard filtered using Quality/average depth estimates to only retain only high confidence variants, which were subsequently used as a known dataset to recalibrate base scores, and 2) final jointly called variants were filtered using the following conservative thresholds (QUAL/Depth > 15, Depth > 5, Mapping Quality > 30, Qual < 10,000, Min. missing data = 25% of samples). After hard filtering, 750,999 variants out of approx. 1.8 million remained. SNP haplotyping was achieved using 10,000 randomly selected SNPS and the program STRUCTURE (V2.3.4) with a burn-in of 20,000, a Y of 20,000, and a K value of 4. To reduce the impact of linkage disequilibrium on inbreeding estimation, the 750,999 variants were thinned by position where no two variants could be located within a very conservative 10kbp. This final filter produced a final variant population of 3066 variants.

The inbreeding coefficient F_*ST*_ was first calculated on a per-sample basis using a method of moments outlined by [33]. As a comparison, F_*ST*_ was also estimated using population delineations according to [34]. Samples were grouped using binary-allelic variant relatedness estimation as outlined by [35] into two populations, North American and European. It was feasible to create two subgroups within the greater European population, United Kingdom samples and mainland Europe, however the resulting F_*ST*_ estimation was not significantly different than the continental comparison and we believe that grouping all European samples is a more conservative hypothesis.

### 1.8 *B. edulis* MAT loci diversity analysis

To asses the theoretical outcrossing efficiency of *B. edulis* we sought to quantify the allelic diversity of the STE3 pheromone receptor. The rcb1.42 STE3-like pheromone receptor gene from *Coprinopsis cinerea* (UNIPROT ID: Q9UVN4-COPCI) was used as an initial reference sequence to extract the approximate gene from BD747 using BLASTn. The exact coding boundaries of the BD747 STE3 gene were determined using previously sequenced transcriptome data from BD747. Seven addition STE3 genes were reconstructed with the same protocol used for ITS reconstruction, from whole genome sequences of two western US samples: 20100815009, DBG24695: and five samples from the southern UK: BD591, BD592, BD593, BD594, BD596. Final reconstructed STE3 sequences were aligned, and the minimum number of novel alleles were counted.

## 2 Results

### 2.1 ITS Diversity

Our initial attempt to reconstruct ITS1-5.8S-ITS2 sequences with traditional bioinformatic methods (method 1 Fig.7) was highly problematic. Initially we attempted to recover ITS sequences from our genome assemblies, however the final ITS consensus sequences were either highly truncated or lacked any polymorphisms representative of their population dynamics. Assembly with Redundans, a genome assembler designed for highly heterozygous genomes, was attempted, yet the short nature of the ITS contig and highly repetitive rDNA array led to failed ITS recovery, except in the instance of the long-short read hybrid assembly of collection BD747, where assembly was partially successful and retained three of the four heterozygous positions. When the ITS region was identified with BLASTn searches using queries from previously assembled genomes sequences, the resulting sequences were truncated on both 5^*1*^ and 3^*1*^ ends. Different assemblers (SPAdes, ABySS, Redundans) produced similar erroneous results. We then aligned raw reads to our *B. edulis* reference and assembled the single ITS contig using ABySS. However, this method produced homogenized sequences from all samples, several of them known to be heterozygotes. To retain any relevant heterozygous information, we manually reconstructed ITS consensus sequences from aligned raw Illumina reads (method 2), and compared these to our data set of 221 *B. edulis* ITS sequences produced from directly-amplified Sanger sequences. We found significantly higher rates of heterozygosity per sample in our genomic ITS sequences compared to our Sanger ITS sequences (1.58 and 0.34 heterozygous positions/sequence respectively; p «0.001). The identity of the heterozygous positions shifted dramatically between the sequence types, from majority N values among the Sanger dataset, where overlapping chromatogram intensities made true identification impossible, to no ambiguous heterozygous positions in the Illumina ITS sequences (Table S2). To identify the approximate proportion of ITS regions presenting each allele in each heterozygous position, we found the number of reads matching each allele and the proportion of reads matching the lower frequency allele/total number of reads. 53.7 percent of all heterozygous positions possess an allele that accounts for less than 40 percent of all reads. This implies that only half of all ITS heterozygous positions in our 28 genomes are potentially the product of recent hybridization events. 14 percent of our heterozygous positions were found at a low frequency, below 5 percent (Table S2). These alleles could be novel mutations in the early stages of rising to fixation via concerted evolution. However, two of the heterozygous positions were present at least twice in the greater *B. edulis* dataset indicating that some of the alleles are the product of previous gene flow or an ancestral metapopulation and are falling to genomic extinction. Moreover, the low-frequency heterozygous position for sample B140 (Table S2) was found to be heterozygous in at least 18 other samples indicating that while rare within the sample, it is not rare within the species. While some of these low-frequency alleles could be novel polymorphisms, at least one is consistent with a previous hybridization event.

**Figure 5:**
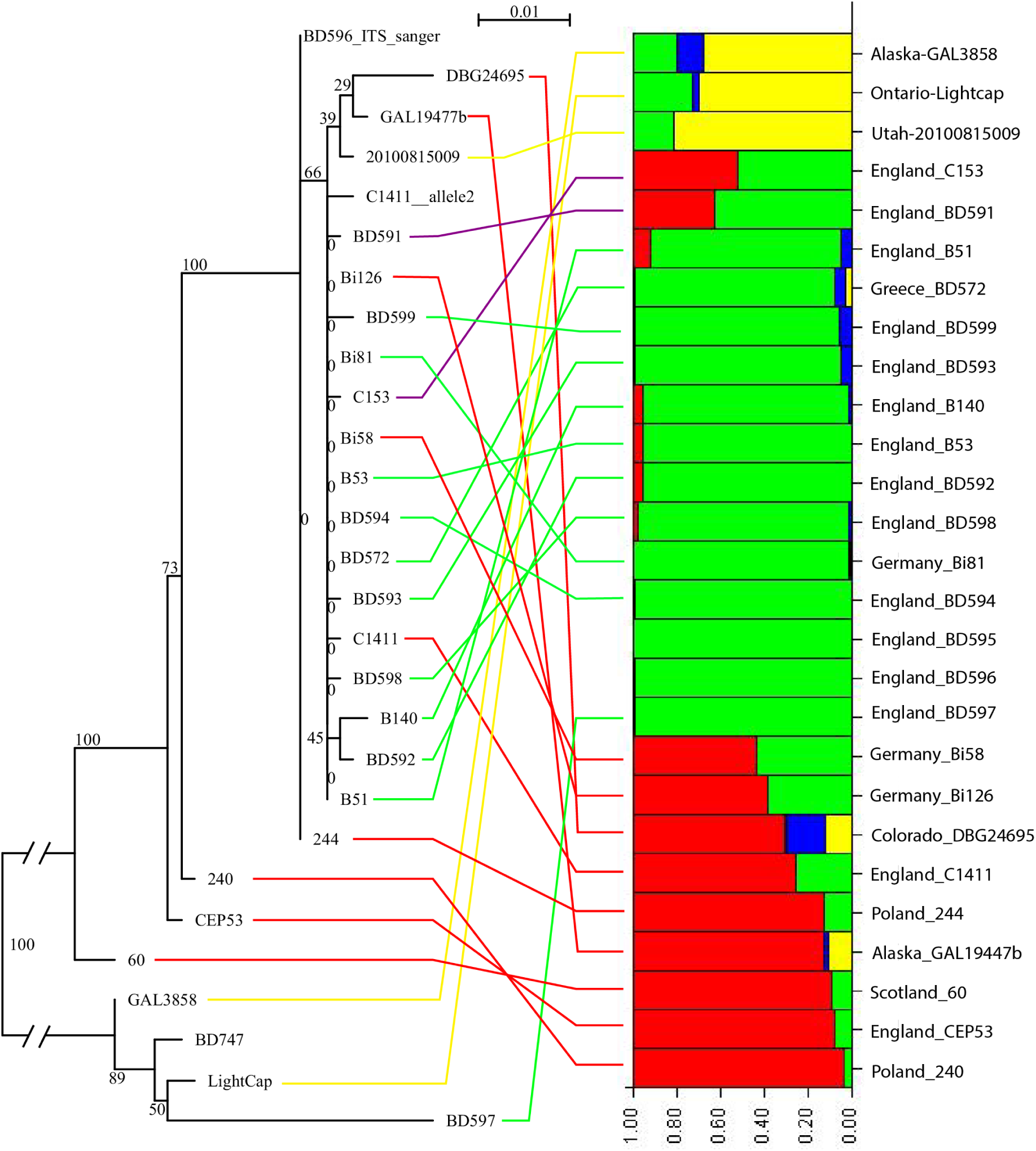
Comparison of ITS gene relatedness using maximum parsimony (left) and whole genome relatedness (right) utilizing 10,000 SNP haplotyping. Size and color of each band in the STRUCTURE plot indicates the proportion of total SNP’s found in that sample aligning to one of the 3 dominant haplotypes. From this comparison we find that ITS barcoding may produce a dishonest signal of relatedness that is not represented throughout the genome.

**Figure 6:**
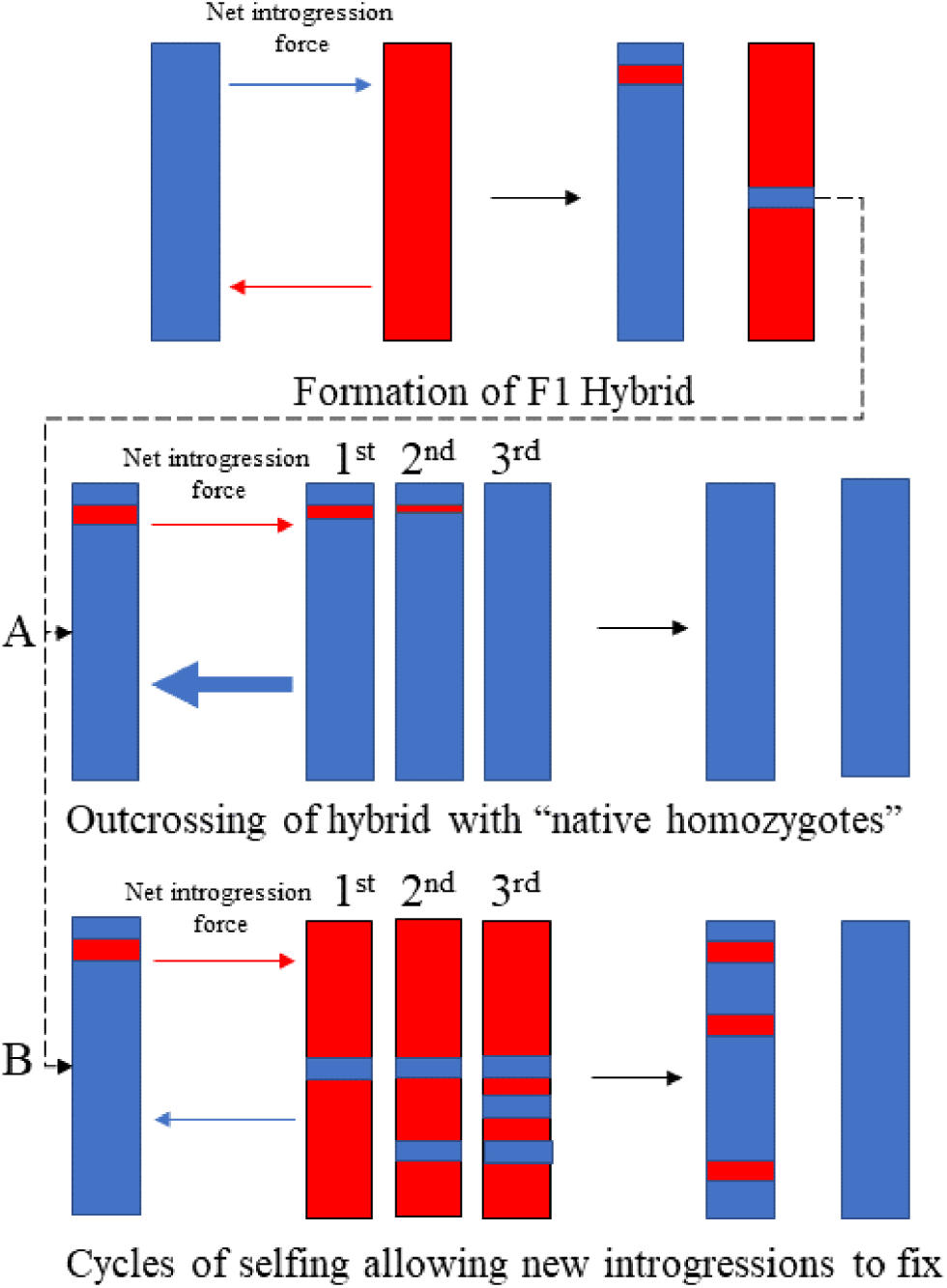
Proposed model of how selfing or inbreeding may produce low frequency allles among the ITS array. After the formation of a F_1_ hybrid, alleles from the “foreign” array (red array) will be stochastically introgressed within the “native” array (blue array). However, since gene flow and hybridization is assumed to be rare, the hybrid “native” array will only interact and recombine with homozygous “native arrays” in subsequent generations (path A). This produces a “net introgression force” that homogenizes the hybrid native array and maintains population ITS homogeneity. However, if inbreeding within populations is high (path B), the likelihood that a hybrid-native array interacts with another hybrid-native or the original foreign array is increased, shifting the balance of net introgression away from homogenization and increasing the likelihood that more “foreign” alleles become introgressed within the native array. With enough generations the new foreign alleles will rise in frequency to the point of fixation within a population, creating a dishonest signal of heterozygosity.

**Figure 7:**
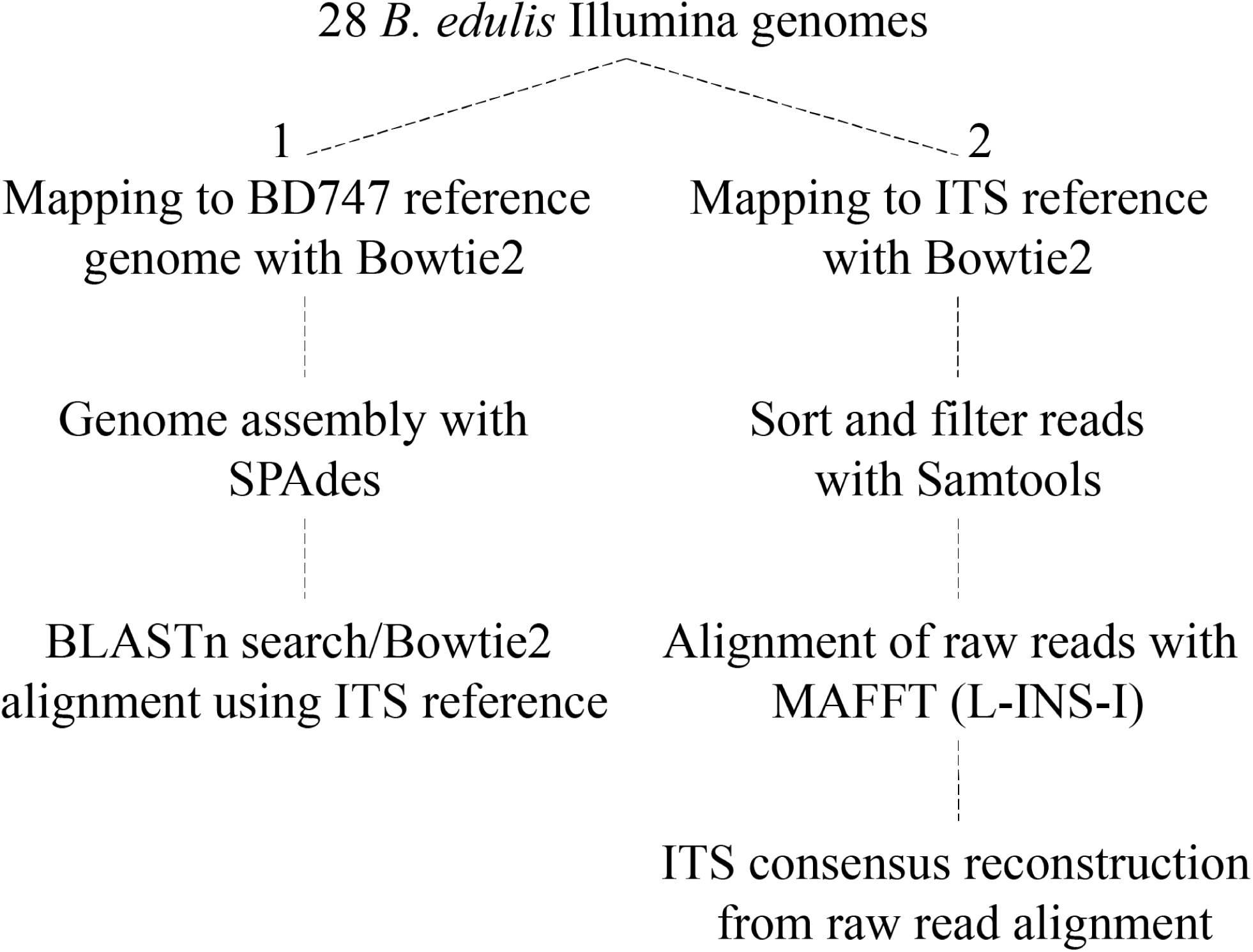
Flowchart of ITS sequence reconstruction using two methods. Method one (left) consists of a stanadard bioinformatic genomic pipeline, while method two (right) was developed to facilitate retention of all polymorphic information along the rDNA array.

### 2.2 Direct comparison of Sanger sequencing and Genomic ITS reconstruction

We were able to compare the ITS sequences of 5 samples obtained from the whole genome and through Sanger sequencing. Of the 14 heterozygous positions found in the genomic ITS sequences, only half were found in the Sanger sequences. Fig. S2 provides an example of a correct and incorrect heterozygote call from Sanger. The average low allele frequency of the positions not called in Sanger sequences was significantly lower than the alleles found in Sanger sequences (mean allele frequency – 0.142, 0.456 respectively p>0.01). While the sample size is limited, our findings indicate that heterozygous alleles that persist at frequencies below approx. 40% of total ITS copies will be lost in standard Sanger ITS barcode sequencing.

### 2.3 Evidence for concerted evolution from Nanopore sequencing

We found evidence of concerted evolution via two of the proposed mechanisms: 1) a recombination event that translocated a large portion of alleles in a homogenous block, and 2) gene conversion that introgressed alleles stochastically from one array to the other (Fig. 4A). From search scheme 1 (see Fig. 4C for a graphical representation of search schemes) we found 121 and 35 reads corresponding to A—A and G—G allele combinations respectively, and no reads corresponding to a mixing of the four alleles. From search scheme 3 utilizing ONT reads, the combination A—A:A—A, which would indicate two ITS regions within the same array was found five times. No reads were found presenting A—A:G—G, indicating that the low frequency alleles (G—G) are not introgressed within larger blocks of majority alleles (A—A). In addition, the hybrid long/short read assembly of BD747 produced three ITS consensus sequences located on four separate scaffolds, which is to be expected from ITS tandem arrays. The majority haplotype A—A was found on both scaffolds while the minority G—G alleles were only found on a single scaffold, alongside the majority combination. Together this provides evidence for a previous recombination event in a F_1_ hybrid individual, where either 1) a small portion of one array underwent recombination and produced a majority/minority heterozygous single array, or 2) recombination produced a true 50/50 heterozygous array and then via concerted evolution one of the two alleles is rising to fixation within the same array. Secondly, from search scheme 2, we searched through raw ONT reads for combinations of the first heterozygous position A/G and the third heterozygous position A/indel 409 bp apart (Table S2). This distance between the two positions was larger than any single Illumina read, necessitating the use of long read technology. We found more than five reads spanning the entire ITS region for every combination of alleles (A—A = 63; A—–indel = 7; G—A = 5; G—indel = 14). Reads spanning two ITS regions (search scheme 3) showed two instances of A—A:G—A, and only four reads presented A—A:A— A. Moreover, the Redundans scaffold assembly of the hybrid (long/short read) BD747 assembly returned both A/indel alleles on two separate scaffolds indicating that no one array is homozygous at the position. Together, this evidence is consistent with concerted evolution via gene conversion, where due to the high sequence similarity between any two rDNA copies, gene conversion will stochastically translocate alleles from one array to its compliment [36] producing an array that has a high diversity of ITS sequences within any one region of the array.

### 2.4 ITS dosage

To estimate the total number of ITS copies present in each genome we calculated the “dosage” by using the average ITS read depth controlled by overall genome depth. Dosage varied widely throughout our dataset (min = 12.619, max = 89.82, mean = 41.475; Fig. S3). Given that background genome coverage also varied between samples we performed a linear regression between genome coverage, and the number of BUSCO’s found in our assembled genomes, and found no significant relationship, indicating that dosage is not a product of sequencing run quality. Unequal recombination has been assumed to rapidly shift rDNA copy numbers as part of concerted evolution and the subsequent shifting of allele frequencies [9, 13, 37]. Intragenomic variation is thought to arise when rDNA copy number is driven to extreme frequencies, therefore we hypothesized that dosage would correlate with ITS heterozygosity. We found no correlation between ITS dosage and average heterozygous allele frequency, total number of heterozygous sites, number of sites <40% in frequency, or the presence or absence of low frequency sites.

### 2.5 Genome-wide Variant Calling, F_*ST*_ Estimation, and haplotype grouping

Across all 28 genomes we initially recovered 750,999 high quality variants from an initial population of 1.8 million. After highly conservative linkage disequilibrium filtering, 3066 variants remained. Using a method of moments analysis that utilizes expected versus observed homozygosity ratios, we estimated individual sample inbreeding, and found high levels across all samples (ratio of 0 = entirely outcrossing, 1 = entirely selfing: min F_*ST*_ = 0.26632, max F_*ST*_ = 0.76612, mean = 0.60936). The three samples with the highest inbreeding coefficients (DBG24695, GAL19447b, GAL3858) are from geographically isolated populations in Colorado and Alaska, providing validating corroboration. In addition, we used a bi-allelic variant assessment of relatedness to parse the sample set into two populations, North American and European, to estimate F_*ST*_ at a higher level. The population level F_*ST*_ also indicated high levels of inbreeding (mean F_*ST*_ = 0.60045).

Haplotype grouping using STRUCTURE and 10,000 randomly selected SNP’s revealed three dominant haplotypes 5. Interestingly, these haplotypes had little geographic structuring. When the haplotypes were compared to an ITS phylogeny produced using maximum prasimony, a strong disconnect between single gene relatedness and whole genome relatedness is found.

### 2.6 *B. edulis* MAT loci diversity analysis

STE3 gene reconstruction from 8 whole genome sequences yielded 65 polymorphic positions across the alignment. DBG24695 possessed three large deletions, the largest 45bp in length, in exon regions. From these 65 polymorphic regions, as a highly conservative assessment, we ascertained that each sample possessed at least one unique STE3 mating-type allele. For the purposes of assessing outcrossing capability, we determined that a unique substitution not found in other samples represented a unique allele. We believe it is far more likely that both alleles possessed by each sample are unique among the population, but without long-read data from all samples it is impossible to link polymorphic sites at the 5^*1*^ and 3^*1*^ ends of the 1370bp gene.

## 3 Discussion

### 3.1 Intragenomic ITS diversity

Analysis of constructed ITS sequences pulled from whole genome sequences has revealed widespread intra-genomic variation in *B. edulis*. Twenty of 28 specimens possessed at least one polymorphic site persisting in a heterozygous state. In addition, this study is the first to directly approximate the number of ITS copies presenting each allele. We found that over 50% of all polymorphic positions persist within individuals at low frequencies among the population of ITS genes. Comparisons of genomic ITS sequences and Sanger ITS sequences highlight the taxonomic and phylogenetic dilemma that these low frequency alleles present. Alleles that persist at below 40% of copies will not be correctly called by Sanger sequencing techniques, yet they may be shared with other individuals in the population, indicating historical or ongoing gene flow. Moreover, from ONT sequencing we have shown that ITS diversity exists within a single ITS array. If the variation persists at a sufficient frequency to be identified by Sanger sequencing, the individual would incorrectly present as a F_1_ hybrid, leading to potentially incorrect taxonomic affinities.

While ITS heterozygosity and intragenomic variation has been previously reported in both plants [38, 39], and *Fungi* [8–10, 40], the rate of variation and heterozygosity may be dramatically under-reported for two reasons: a) These studies have relied on sequence identification using RFLP-gel electrophoresis or Sanger Sequencing analysis, which we have shown will underreport variation due to the presence of low-frequency heterozygosity, and b) vector-cloning PCR used in these studies is agnostic with regards to the allelic structure of the rDNA arrays. Altogether, this study provides evidence for unprecedented intra-specific, intra-genomic, and intra-array variation. In addition, we highlight the limitation of Sanger sequencing for identification of heterozygosity or low frequency ITS polymorphisms. Naidoo et al. [5] found evidence for rapid change in allelic composition during a single round of meiotic division. They proposed that unequal recombination (UR) was the primary force behind such dramatic change in allelic frequency and gene conversion can lead to allele fixation after subsequent rounds of recombination. UR within the rDNA array has been thought to stochastically shift cistron copy numbers [1, 5]. ITS dosage varied widely among our samples, from approx. 13 to 90(Fig. S3), highlighting the intraspecific diversity in rDNA copy number in *B. edulis*, which provides further evidence that UR is an active and dynamic force among the ITS array. However, UR alone may not be sufficient to drive allele frequencies to low frequency as suggested by Naidoo et al. [5]. Based on their model, after the formation of an F_1_ hybrid, UR will increase or decrease rDNA copy number during primarily meiosis and drive one allele to high or low frequency. We found no correlation between ITS dosage and average heterozygous allele frequency, total number of heterozygous sites, number of sites below 40% in frequency, or the presence or absence of low frequency sites. This suggests that the presence or absence of low frequency alleles in the ITS is not directly associated with copy number, suggesting that gene conversion is a more active force than previously thought. Further evidence from ONT sequencing has shown that low frequency alleles can exist within the ITS population as a single block within one array, and introgressed in all combinations within both ITS arrays. The existence of a single low-frequency homogeneous allele block within a single array is most likely the product of an UR event. In contrast, ITS alleles that are mixed in all combinations is more likely the product of stochastic gene conversion and the subsequent introgression of alleles from both parental arrays. Direct evidence supporting gene conversion as a mechanism of concerted evolution has unfortunately been rare [1]. However, here we have potentially presented the first direct evidence of gene conversion in basidiomycete *Fungi* involved in homogenizing the rDNA array.

The presence of intragenomic variation in the ITS array has been thought to be due to the relaxation of concerted evolution. However, the introgression of alleles from one parental ITS array into the compliment via both recombination and gene conversion indicates that the proposed mechanisms of concerted evolution [2] are still active in populations of *B. edulis*. The relative timing for homogenization via converted evolution has conflicting experimental evidence. Fuertes-Aguilar et al [41] found that F_1_ hybrids of two species of *Armeria* (Plumbaginaceae) presented true Hardy-Weinburg heterozygotes in ITS, yet were homozygotes by the third generation. In contrast, *Nicotiana* allopolyploids may take hundreds to thousands of years to homogenize rDNA arrays [39]. However, these studies in plants were limited to identification of interarray variation and similar patterns have never been analyzed in basidiomycetes. How ITS variation is introgressed from one array to another and the latency period of this variation is entirely unknown. We propose that subsequent cycles of inbreeding or selfing after the formation of a hybrid between two populations would sufficiently fix introgressed alleles within an array (Fig. 4). Within an F_1_ hybrid, a small number of ITS genes will stochastically be transferred via concerted evolution from the “foreign” parental ITS array into the “native” array. However, given that populations of *B. edulis* exist over large spatial distances in North America [42], form large conspicuous fruiting bodies, yet colonize their hosts at low rates [43], long distance dispersal and gene flow between any two populations must be exceedingly rare, and therefore, the likelihood that an F_1_ hybrid mates with another F_1_ hybrid is equally rare. Outcrossing of the F_1_ hybrid and “native” ITS homozygotes will create a net homogenizing force of gene conversion that will eliminate the newly introgressed ITS alleles in the F_1_ hybrid (Fig. 6). Yet if inbreeding and selfing play a larger role in *B. edulis* than previously thought, the F_1_ hybrid will then potentially undergo several mating cycles between the “native” and “foreign” ITS arrays, allowing introgressed alleles from the “invader” to rise to sufficient fixation frequency in the “native” array. After fixation, the newly introgressed alleles will persist for several generations at a low frequency within the array or rise to complete fixation within the population. This hypothesis is supported by our results in three ways: 1) high levels of inbreeding were estimated for all samples, 2) highly varied ITS dosage indicates rampant UR in samples, and 3) introgression of alleles creates variation within chromatids. Furthermore, reproductive isolation by distance and strong population structuring have been found in several other basidiomycete taxa [44], suggesting this pattern may be the rule rather than the exception. For example, sequencing of the intergenic spacer (IGS) region in *Tricholoma scalpturatum* found strong fine-scale genetic spatial autocorrelation, which highlights the inefficacy of spore dispersal [45].

### 3.2 Inbreeding persists despite high theoretical outcrossing efficiency

To verify that the high inbreeding rates found in *B. edulis* are due to patterns of gene flow and not the product of genomic constraints, such as low MAT allelic diversity or the loss of a MAT locus, we calculated the approximate allelic diversity of the STE3 pheromone receptor. Importantly, we found that, regardless of geographic proximity, each specimen possessed at least one novel STE3 allele, and that the number of unique alleles is equal to the number of specimens sampled. This diversity is equivalent with the diversity found in other basidiomycete taxa that have high theoretical outcrossing efficiency [46]. In addition, gene annotation of JGI’s *Boletus edulis* BED1 v4.0 indicates the existence of the second MAT-locus encoding a homeodomain transcription factor, confirming that *B. edulis* can be classified as possessing a tetrapolar mating system. This indicates that *B. edulis* has a high theoretical outcrossing efficiency and gene flow is not constrained by low allelic diversity at the mating type loci. However, our samples are nonetheless highly inbred, perhaps highlighting a disconnect between the ability to sexually reproduce and true gene flow between populations and individuals.

### 3.3 Heterozygosity in ITS may not be indicative of actual gene flow

Heterozygous positions in ITS have long been thought to be a direct indication of gene flow between populations [15]. Our samples have on average 1.58 heterozygous positions per sequence, which would suggest rampant gene flow between distant populations. Contradicting this observation, genome-wide fixation indices are consistent with highly inbred populations, indicating relative low amounts of gene flow. Moreover, using ONT sequences, we document that some of these heterozygous positions are stochastically introgressed among and between ITS arrays (Fig. 4A) and are therefore not F_1_ hybrids at these positions. We believe that inbreeding and concerted evolution are artificially highlighting the signal of rare gene flow events in the ITS array that is not consistent with organismal recombination rates at the population level.

## 4 Conclusions and implications for *B. edulis* and other taxonomic inferences

If we were to asses our ITS dataset from only Sanger sequences without corroborating single copy genes, it is likely that the highly structured populations and ITS variation would be interpreted as populations in the process of speciating or having recently diverged. This may even lead to spurious taxonomic assignment, such as recognizing this population structure as distinct species. However, when new sequence information is considered using genomic ITS reconstruction, we find that population clustering decreases and hybridization events between populations are common. While the presence of ITS variation in and of itself may indicate cryptic speciation, we believe that it is a dishonest signal, and instead propose that the lifestyle characteristics of *B. edulis* may be artificially elevating the presence of intra-specific variation via the mechanisms of concerted evolution. In summary, significant intraspecific variation in ITS sequences is not sufficient to indicate cryptic speciation events, the mechanisms of concerted evolution may be insufficient to homogenize the rDNA array in some species, and the natural history of a species can complicate the use of ITS barcoding for taxonomic quantification and identification. While the routine use of ITS barcodes for distinguishing well-definied species is not compromised by these results, its use for resolving population/species boundaries or for delineating cryptic species is inappropriate without corroborating evidence from multilocus sequence data or other information.

## Acknowledgments

We are grateful to Bill Amos, Joe Ammirati, Tim Baroni, Brad Kropp, Mary Smiley, and Igor Safanov, as well as the Burke Museum, for providing specimens used in this study. We would like to thank Nathan Smith for his effort in sequencing and data production. Thanks to Phil Madgwick for help with lab work and Sanger sequence editing. Funding was provided in part by the Charles Wolfson Trust (London, UK).

