## Supplemental Figure 2 for "Lost in Translation: population genomics and long-read sequencing reveals relaxation of concerted evolution of the ribosomal DNA cistron"

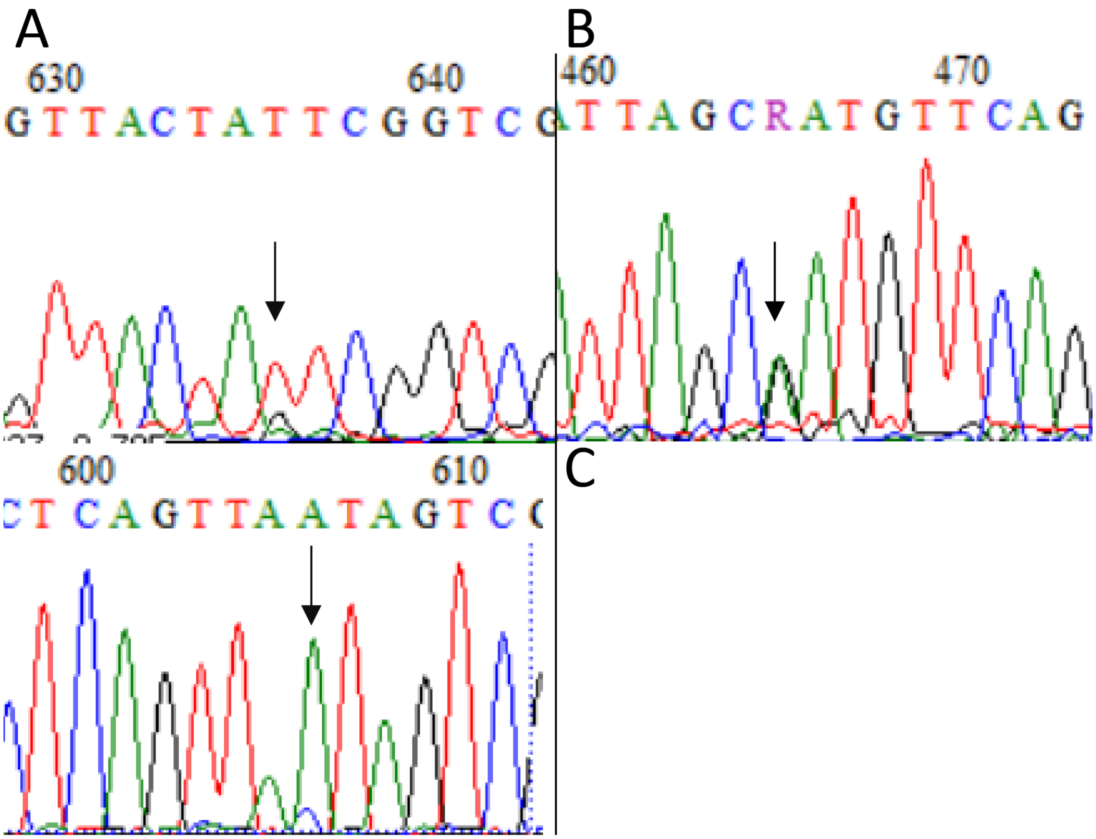


Fig. S2. Three electropherograms covering heterozygous positions indicated by black arrows. Panels A and C represent heterozygous positions (T/G and A/C respectively) where one allele exists at a low frequency among ITS copies. Panel B indicates a heterozygous position (A/G) in near hardy-Weinberg proportions. While the low frequency allele is present as a small hump, the intensity is low enough that it will not be automatically called and appears as background noise.
