## Supplemental Figure 3 for "Lost in Translation: population genomics and long-read sequencing reveals relaxation of concerted evolution of the ribosomal DNA cistron"

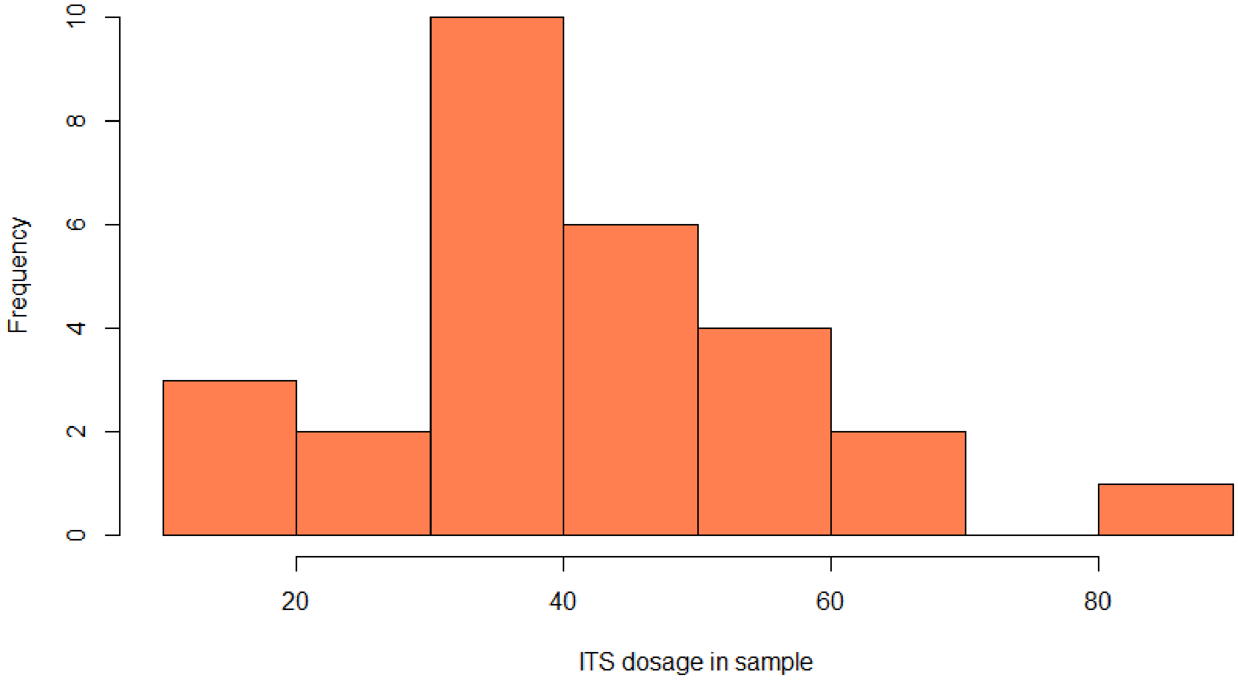


Fig. S3. Histogram of ITS dosage found in 29 whole genome sequences of *B. edulis* . While not an exact count of rDNA copy number in each specimen, dosage allows us to approximate the relative diversity of copy number among our population. We found that dosage varied dramatically, from 13 to 90 indicating that unequal recombination is active in our samples.
