## Supplemental Figure 1 for "Lost in Translation: population genomics and long-read sequencing reveals relaxation of concerted evolution of the ribosomal DNA cistron"

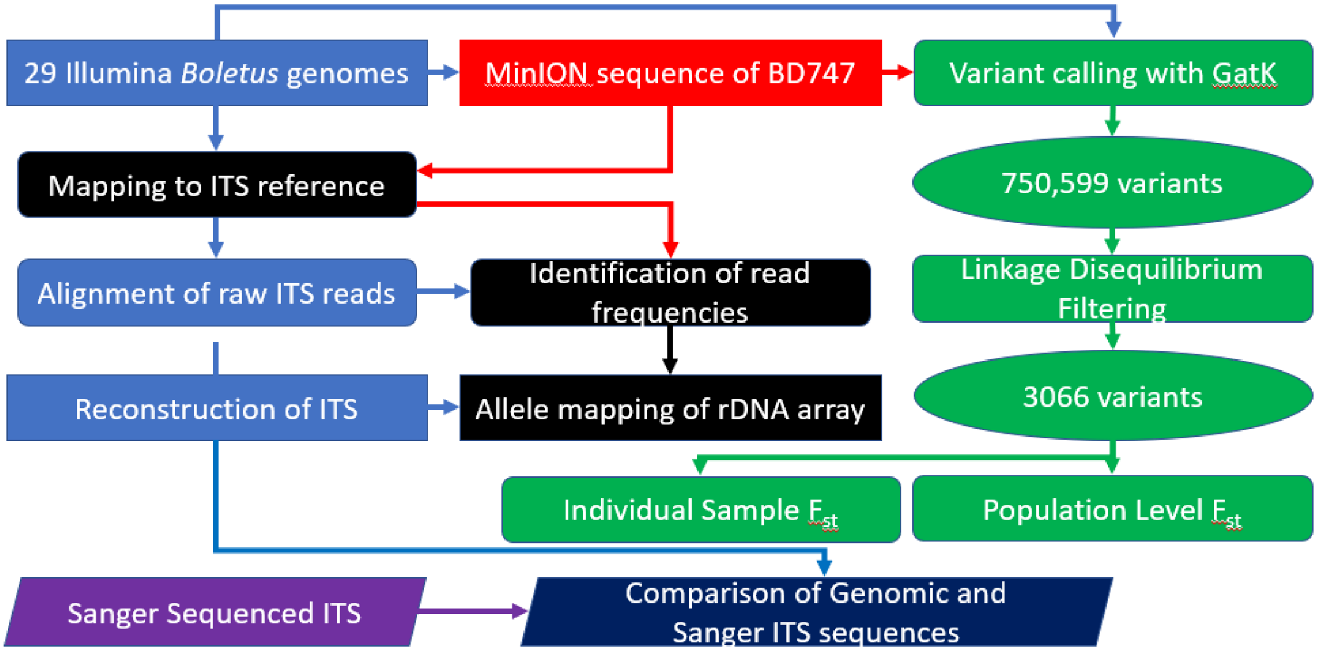


Fig. S1. Flowchart of bioinformatic pipelines used in this study. Color of box is indicative of sequencing data used for analysis. For example blue indicates Illumina short reads and red indicates MinION long reads.
